# Identification of Functional Noncoding RNA-encoded Proteins on Lipid Droplets

**DOI:** 10.1101/2020.04.10.036160

**Authors:** Ting Huang, Adekunle T. Bamigbade, Shimeng Xu, Yaqin Deng, Kang Xie, Ololade O. Ogunsade, Ahmed Hammad Mirza, Jifeng Wang, Pingsheng Liu, Shuyan Zhang

## Abstract

Over the past decade, great progress in sequencing technologies and computational biology has revealed that the majority of the mammalian genome considered to be noncoding is rich in functional elements able to produce proteins. Many RNA molecules, mis-annotated as noncoding, actually harbor small open reading frames that are predicted to code for proteins. Some of those proteins have been verified to play critical roles in multiple biological processes. The lipid droplet (LD) is a unique cellular organelle, conserved from bacteria to humans, and is closely associated with cellular lipid metabolism and metabolic disorders. No noncoding RNA-coded proteins have been identified on LDs. Here, for the first time, we searched the organelle for their presence. After the enrichment of small proteins of LDs isolated from myoblasts, we used mass spectrometry coupled with our lab made protein database to identify LD-associated noncoding RNA-encoded proteins (LDANPs). A total of 15 new proteins were identified. One of them was studied further and termed LDANP1. LDANP1 was localized on LDs by imaging, cell fractionation, and immunogold labeling. Like LD resident proteins, LDANP1 was degraded by the proteasome. Using the CRISPR/Cas9-mediated genome editing technique, the endogenous expression of LDANP1 was validated. The stable expression of LDANP1 suppressed the accumulation of triacylglycerol in oleic acid treated myoblasts and inhibited the rescue of palmitate-inhibited insulin sensitivity by oleic acid. In summary, we report for the first time that translatable, nominally noncoding RNA-derived proteins, which are new and cannot be identified using current research methods, were associated with LDs and that among these, LDANP1 modulated lipid metabolism and insulin sensitivity. The discovery of noncoding RNA-encoded proteins on LDs paves a new way for the research of LDs and lipid metabolism.

## 1. Introduction

It has been reported for years that only 1.5% of the sequence of the human genome actually contains instructions for making proteins[1]. However, in recent years with the rapid development of high-throughput sequencing technologies, it has become evident that the eukaryotic genome is extensively transcribed[1–4]. Large-scale projects for the systematic annotation and functional characterization of genes, such as GENCODE and FANTOM, demonstrate that at least 80% of mammalian genomic DNA is actively transcribed. This has led to the identification of numerous nonclassical protein-coding genes, which were previously thought to be noncoding sequences[5–7]. It has become increasingly clear that many genomic regions classified as noncoding harbor short open reading frames (sORFs) with fewer than 300 nucleotides, coding for small, functional proteins or micropeptides. The translated products have fewer than 100 amino acids which has historically been the minimum length for detection. These small proteins have evaded detection due to their small size, low abundance, instability, and loss during sample preparation[7]. However, emerging experimental techniques, including ribosome profiling and mass spectrometry, and computational advances have permitted discovery and analysis of those proteins[7; 8]. Most micropeptides have been identified in prokaryotic organisms[8] but a growing handful have also been discovered in eukaryotic organisms, including plant, zebra fish, insect, mouse, and humans[4; 7; 8]. Micropeptides have been found to play important roles in many fundamental biological processes. For instance, the micropeptide Toddler has been proven to be an embryonic signal that promotes cell movement in zebrafish[9]. Myoregulin (MLN) co-localizes with and inhibits SERCA, a Ca2^+^ pump found in the sarcoplasmic reticulum of skeletal muscle, and thus affects calcium homeostasis in mice[10]. MOTS-c, a mitochondrial-derived peptide, is reported to promote metabolic homeostasis and to reduce diet-induced obesity and insulin resistance in mice[11]. Proteins from nominally noncoding transcripts have been reported to localize to some cellular compartments such as mitochondrion[11], late endosome/lysosome[12], and endoplasmic reticulum[10]. However, given the large number of putative proteins encoded by noncoding-RNAs, much more investigation is required. In particular, more information is needed on the subcellular localization of these proteins, which may provide insight into their biological functions.

The lipid droplet (LD) is a ubiquitous intracellular organelle containing a neutral lipid core bounded by phospholipid monolayer membrane and some integral and peripheral membrane proteins[13; 14]. LDs play critical roles in lipid storage, transport, and metabolism[15–17]. Abnormal LD dynamics is associated with the development of metabolic diseases, such as obesity, atherosclerosis, fatty liver disease, and type 2 diabetes[17–19]. The LD-associated proteins are the main executors of LD function. For instance, Perilipin 2 (PLIN2), the most abundant LD resident protein in cells except for adipocytes, has been proven to function in hepatic lipid sequestration[20] and improves insulin sensitivity in skeletal muscle[21]. The most prevalent proteins reported to be associated with LDs are enzymes involved in lipid hydrolysis and synthesis[17]. Continued study of LD proteins is crucial to understand the role of LDs in normal homeostasis and metabolic diseases. Over 100 proteins have been found on LDs and many of them have been associated with human metabolic diseases[17; 18]. However, there have been no reports identifying LD-associated proteins derived from nominally noncoding transcripts.

Here for the first time, the existence of noncoding RNA-encoded proteins on LDs was explored. We adopted an experimental approach to identify LD-associated noncoding RNA-encoded proteins (LDANPs) in myoblast cells. LDs were isolated and the enriched small proteins less than 17 kDa were subjected to mass spectrometric study. A home-made database that we setup previously for identification of noncoding RNA-encoded proteins was used and a total of 15 novel proteins derived from noncoding RNAs were identified in C2C12 LDs. The top one LDANP, LDANP1, was further studied in detail. This novel protein was shown to localize on LDs and degraded though proteasome. When loaded with oleate (OA), LDANP1-overexpressing cells displayed a significant reduction in triacylglycerol (TAG) and rescue of palmitate (PA)-reduced insulin sensitivity was inhibited.

## 2. Materials and methods

### 2.1. Materials

The Colloidal Blue Staining Kit, Hoechst 33342 and LipidTOX Red were from Invitrogen. Sodium oleate and sodium palmitate were obtained from Sigma-Aldrich. Western lightning plus-ECL reagent was from PerkinElmer. 8% paraformaldehyde solution (EM grade) and LR White resin were purchased from Electron Microscopy Sciences. The synthesized PCR primers used are listed in Supplemental Table S2. Antibodies used in this study are listed in Supplemental Table S3.

### 2.2. Cell culture

Mouse C2C12 myoblasts (American Type Culture Collections, Manassas, VA) were maintained in DMEM (Macgene Biotech., Beijing) supplemented with 10% FBS (Hyclone), 100 U/mL penicillin and 100 μg/mL streptomycin at 37°C with 5% CO_2_.

### 2.3. Preparation of fatty acid

Sodium oleate and sodium palmitate were prepared according to our previously established method[22]. Briefly, fatty acids were mixed with ethanol to a concentration of 100 mM and sonicated on ice at 200 W, 10 s on and 3 s off pulses until the solution was milky and homogenous. The prepared fatty acid stocks were kept at 4°C and kept from light. The fatty acid was incubated in 60°C preheated medium before treatment of cells.

### 2.4. Isolation of lipid droplets from C2C12 cells

LDs were isolated from C2C12 cells using the method described previously with modifications[23; 24]. Briefly, C2C12 myoblasts treated with 100 µM OA for 12 h were scraped and collected after 3 rinses with ice-cold PBS. Then all the cells were transferred to 50 mL Buffer A (25 mM tricine, 250 mM sucrose, pH 7.8) containing 0.5 mM PMSF. After centrifugation at 3,000*g* for 10 min the cell pellets were re-suspended in 10 mL Buffer A containing 0.5 mM PMSF and incubated on ice for 20 min. The cells were then homogenized by N_2_ bomb (750 psi for 15 min on ice). The cell lysate was centrifuged at 3,000*g* for 10 min, the post-nuclear supernatant (PNS) fraction was collected and loaded into a SW40 tube and the sample was overlain with 2 mL Buffer B (20 mM HEPES, 100 mM KCl, and 2 mM MgCl_2_, pH 7.4). The sample was centrifuged at 182,000*g* for 1 h at 4°C. The white band containing LDs at the top of gradient was collected into a 1.5 mL Eppendorf tube. The LD sample was centrifuged at 20,000*g* for 10 min at 4°C and then the underlying solution was carefully removed. The LDs were gently re-suspended in 200 μL Buffer B. This procedure was repeated four times. The lipid extraction and the protein precipitation were carried out using chloroform/acetone (1:1, v/v) treatment followed by centrifugation at 20,000*g* for 10 min at 4°C. The protein pellet was then dissolved in 2 × SDS sample buffer (125 mM Tris Base, 20% glycerol, 4% SDS, 4% β-mercaptoethanol and 0.04% bromophenol blue, pH 6.8).

### 2.5. Western blot analysis

Whole cell lysates were suspended in 2 × sample buffer, sonicated, and denatured for 5 min at 95°C. 10 µL of sample was subjected to gel electrophoresis followed by silver staining or protein blotting onto 0.2 µm PVDF membrane. To detect proteins of interest, membranes were incubated with the indicated antibodies and were detected using ECL.

### 2.6. Immunoprecipitation

Cells were cultured to 100% confluence and the culture medium was removed by aspiration. Cells were washed with cold PBS three times and were then suspended in 1 mL cold TETN (25 mM Tris-HCl, 5 mM EDTA, 1% Triton X-100, pH 7.5) solution containing 150 mM NaCl supplemented with 0.5 mM PMSF. The mixture was left on ice for 30 min. Cells were scraped into a protease-free Eppendorf tube and pipetted 8 times followed by rotation at 4°C for 30 min. In a separate tube, FLAG beads (SIGMA M2 Affinity Gel) were blocked with 20 mg/mL BSA in TETN solution containing 150 mM NaCl. Then, the beads were rotated at 4°C for 10 min, followed by centrifugation at 1,000*g* for 30 s. The supernatant was removed, and this process was repeated three times. After 30 min rotation, the whole cell lysate was centrifuged at 17,000*g* for 10 min at 4°C. An aliquot of the supernatant was kept as ‘Input’ and was mixed with an equal volume of 2 × sample buffer. The remaining supernatant was incubated with FLAG beads, previously blocked with BSA, at 4°C for 3 h. The supernatant-bead complex was centrifuged at 1,000*g* for 30 s at 4°C. An aliquot of the supernatant was kept as ‘Supernatant’ while the remainder was discarded. The beads were washed with TETN containing 500 mM NaCl followed by TETN containing 250 mM NaCl and finally with TETN containing 150 mM NaCl, twice for each buffer. The supernatants were discarded, and the beads were suspended in 50 µL 2 × sample buffer, vortexed vigorously for 5 min, and denatured at 95°C for 5 min (IP sample).

### 2.7. LC-MS/MS Analysis

The LD proteins were separated on a 12% Bis-Tris gel with a 4% stacking gel. The gel was then stained with Colloidal Blue according to the manufacturer’s protocol (Invitrogen). The protein band less than 17 kDa was cut into slices and the protein sample was manually excised, cut into small plugs, and washed twice in 200 μL of ddH_2_O for 10 min each time. The gel bands were dehydrated in 100% acetonitrile for 10 min and dried in a Speedvac for approximately 15 min. Reduction (10 mM DTT in 25 mM NH_4_HCO_3_ for 45 min at 56°C) and alkylation (40 mM iodoacetamide in 25 mM NH_4_HCO_3_ for 45 min at room temperature in the dark) were performed, followed by two washes of the gel plugs with 50% acetonitrile in 25 mM ammonium bicarbonate. The gel plugs were then dried using a Speedvac and were digested with sequence-grade modified trypsin (40 ng for each band) in 25 mM NH_4_HCO_3_ overnight at 37°C. The enzymatic reaction was stopped by the addition of formic acid to a 1% final concentration. The solution was then transferred to a sample vial for LC-MS/MS analysis. Besides, half of the extracted LD proteins were not stained after electrophoresis and cut directly for proteomic study.

All nano LC-MS/MS experiments were performed on a Q Exactive (Thermo Scientific) equipped with an Easyn-LC 1000 HPLC system (Thermo Scientific). The peptides were loaded onto a 100-μm id × 2-cm fused silica trap column packed in-house with reversed phase silica (Reprosil-Pur C18-AQ, 5 μm, Dr. Maisch GmbH) and then separated on an a 75-μm id × 20-cm C18 column packed with reversed phase silica (Reprosil-Pur C18-AQ, 3 μm, Dr. Maisch GmbH). The peptides bound on the column were eluted with a 78-min linear gradient. The solvent A consisted of 0.1% formic acid in water and the solvent B consisted of 0.1% formic acid in acetonitrile. The segmented gradient was 4–12% B, 5 min; 12–22% B, 53 min; 22–32% B, 12 min; 32-90% B, 1 min; 90% B, 7 min at a flow rate of 300 nL/min.

The MS analysis was performed with Q Exactive mass spectrometer (Thermo Scientific). In a data-dependent acquisition mode, the MS data were acquired at a high resolution 70,000 (*m/z* 200) across the mass range of 300–1,600 *m/z*. The target value was 3.00E+06 with a maximum injection time of 60 ms. The top 20 precursor ions were selected from each MS full scan with isolation width of 2 *m/z* for fragmentation in the HCD collision cell with normalized collision energy of 27%. Subsequently, MS/MS spectra were acquired at resolution 17,500 at *m/z* 200. The target value was 5.00E+04 with a maximum injection time of 80 ms. The dynamic exclusion time was 40 s. For nano electrospray ion source setting, the spray voltage was 2.0 kV; no sheath gas flow; the heated capillary temperature was 320°C.

### 2.8. Protein identification

The raw data from the Q Exactive were analyzed with Proteome Discovery (version 1.4.1.14, Thermo Scientific) using Sequest HT search engine for protein identification and Percolator for FDR (false discovery rate) analysis against our custom protein database constructed from noncoding transcripts[25]. Some important searching parameters were set as follows: two missed cleavages were allowed for searching; the mass tolerance of precursor was set as 10 ppm and the product ions tolerance was 0.02 Da.; the methionine oxidation was selected as variable modifications; the cysteine carbamidomethylation was selected as a fixed modification. FDR analysis was performed with Percolator and FDR <1% was set for protein identification. The peptides confidence was set as high for peptide filter. For the stained sample, the raw MS data was searched with parameter of trypsin selected as enzyme as well as no-enzyme as parameter while for unstained sample only using no-enzyme parameter.

### 2.9. Plasmid construction

The LDANP1 ORF was amplified from exon 4 in the USPL1 (UniProtKB entry Q3ULM6) coding sequence. LDANP1 was cloned into pEGFP-N1 with XhoI and HindIII restriction endonucleases. The recombinant plasmid (LDANP1-GFP cDNA) was in turn amplified with a primer pair containing FLAG epitope in the reverse primer and was cloned into pQCXIP with NotI and EcoRI restriction endonucleases where GFP-FLAG served as control. To change Lys to Ala in LDANP1, point mutations at positions 7, 49, 61 and 80 were carried out using the QuikChange mutagenesis system (Agilent) according to manufacturer’s protocol. Mutant LDANP1-GFP fusion proteins were subcloned into pQCXIP plasmid. List of primers is provided in Supplemental Table S2.

### 2.10. Constitutive expression of LDANP1 in C2C12 Cells

C2C12 cells stably expressing LDANP1were constructed using Clontech retroviral packaging system. For virus packaging, VSVG (vesicular stomatitis virus G) envelop plasmid and pHIT packaging plasmid were mixed together in Lipofectamine 2000 according to instruction manual along with LDANP1-GFP-FLAG recombinant gene in pQCXIP plasmid. The plasmid mix was transfected into Plat-ET cells. Medium containing the packaged virus was filtered with 0.45-µm filter to remove dead cells. Wild type (WT) C2C12 cells were cultured in 6-well dishes to a confluence of 40% and were infected with medium containing LDANP1 expressing virus supplemented with 8 μg/mL polybrene to increase infection efficiency. 48 h post-infection the cells were sub-cultured into 100-mm dish and were selected with 1 µg/mL puromycin for two weeks. Immediately after the selection process, cells were dispensed into 96-well dishes to select for clones.

### 2.11. CRISPR/Cas9-mediated genome editing of LDANP1

To construct homologous arms for LDANP1, the nucleotide sequence spanning the region 1,000 bp upstream to 1,000 bp downstream of the reference stop codon was amplified and assembled in pKI-FLAG-N1 incorporating the 3×FLAG as previously established[26]. The 3×FLAG was encrypted in a 34-codon sequence with an additional linker sequence (7 codons). CRISPR/Cas9 technique was employed to induce double strand breaks in the target genes in the C2C12 cell genome. The sequence of the last exon/intron that includes approximately 50 bp upstream or downstream of the stop codon on the left and right was analyzed using the interactive webpage http://crispr.mit.edu/ to design the sgRNA most suitable to induce double strand break in the genome. A short sequence containing 20 nucleotides was synthesized with CACC and AAAC flanking the forward and reverse primers, respectively, in the 5’ directions. The primers were annealed and ligated into a px260a plasmid previously restricted with BbsI.

Knock in plasmids were co-transfected with the respective sgRNA plasmids into C2C12 myoblast by electroporation. Forty-eight hours (48 h) post-transfection, cells were selected with 1 µg/mL puromycin for two weeks with a medium change every 48h. Surviving cells were screened for clones expressing FLAG fusion protein both by genotyping and Western blot.

### 2.12. Fluorescence microscopy

C2C12 myoblasts were transfected with 0.5 µg LDANP1 with C-terminally fused GFP (pEGFP-N1). Transfected cells were cultured on glass cover slips in a 24-well dish with empty plasmid as a transfection control (pEGFP-N1). Cells were treated with OA-containing medium for 12 h prior to sample preparation for image capture. Twenty-four hours (24 h) post-transfection, cells were rinsed with cold PBS three times and stained with LipidTOX Red (1:1,000) for 30 min, and Hoechst 33342 (1:1,000) for 10 min. Cells were rinsed three times with PBS and the cover slips were placed on confocal dish for image capture using Olympus FV1200 confocal microscope (Olympus Corp, Lake Success, NY).

### 2.13. Fluorescence immunostaining

After incubation with OA for 12 h, C2C12 cells stably expressing GFP-FLAG and LDANP1-GFP-FLAG grown on glass cover slips were rinsed with cold PBS three times, fixed with 4% (v/v) paraformaldehyde for 30 min, and then 50 mM NH_4_Cl in PBS to quench residual aldehydes. The cells were then permeabilized with 0.1% Triton X-100 for 15 min. 3% BSA was used to block non-specific antibody binding. Cells were then incubated with primary antibody (anti-GFP or anti-FLAG, 1:100) for 2 h and subsequently with secondary Alexa Fluor 488-conjugated antibody for 1 h. The cells were further stained with LipidTOX Red (1:1,000) for 30 min, and Hoechst 33342 (1:1,000) for 10 min. Images were captured with Olympus FV1200 confocal microscope.

### 2.14. Immunogold labeling

The localization of LDANP1 was visualized by immunoelectron microscopy. In brief, cells stably expressing LDANP1-GFP-FLAG were treated with 100 µM OA plus 1 µM MG132 for 12 h. Cells were collected by trypsin digestion and were fixed in 4% (v/v) paraformaldehyde overnight at 4°C. The cells were subsequently dehydrated in an ascending ethanol concentration series at room temperature, including 50%, 70%, and 80%, for 15 min each. The samples were incubated in a 2:1 mixture of LR White resin to 70% ethanol for 1 h, and in three changes of 100% LR White, for 1 h, 12 h and 1 h respectively. Cells in LR White resin were placed in a gelatin capsule to polymerize at 50 °C for 24 h. The resin block with cells were cut into 70 nm sections on Leica EM UC6 Ultramicrotome. The ultrathin sections were mounted on a formvar-coated copper grid. For labeling, the grids were incubated on drop of PBS with 1% BSA and 0.15% glycine for 10 min, then transferred to drop of GFP polyclonal antibody (TaKaRa, Cat. No. 632592) at 1:50 for 1 h and subsequently in second antibody conjugated with 18-nm gold particles (Jackson, Cat. No. 111-215-144) at 1:100 for 1 h. After being stained with 1% uranyl acetate for 10 min, the sections were observed with Tecnai Spirit electron microscope (FEI, Netherlands).

### 2.15. Total cellular triacylglycerol quantitation

C2C12 cells were cultured in 6-well dishes to 75% confluence. Subsequently, cells were incubated in DMEM supplemented with or without 100 µM OA for 12 h. Cells were washed with PBS three times by using 2 mL PBS each time. To each well, 0.3 mL of 1% (v/v) Triton X-100 in PBS was dispensed and incubated at 37°C for 10 min. Cells were collected and sonicated using 200 W, a 9 s pulse on, 4 s pulse off setting for 5 times. Ten microliter (10 µL) volume of whole cell lysate was used for triacylglycerol analysis with total protein serving as an internal control. Equal aliquots of whole cell lysate were used for protein analysis. Briefly, protein standard supplied with BCA (Bicinchoninic acid Assay) Protein Kit was diluted according to the manufacturer’s instructions with a maximum concentration of 2 mg/mL. Each concentration ranging from 0 mg/mL to 2 mg/mL was prepared in triplicate. Ten microliters (10 µL) of each standard concentration was dispensed into a 96-well dish and 190 µL of BCA chromogenic agent was added. For samples of unknown protein concentration, a similar procedure was followed. Samples were incubated at 37ºC for 45 min and the optical density (OD) was read at 562 nm. Following a similar procedure, whole cell lysate samples were analyzed for triacylglycerol, using appropriate standards and chromogenic agent. Triacylglycerol standard ranged from 0 mg/dL to 40 mg/dL. Upon addition of chromogenic agent mix to sample, assay mixture was incubated at 37ºC for 10 min and the OD was read at 505 nm.

### 2.16. Statistical analyses

Data were presented as mean ± SEM. The statistical analyses were performed using GraphPad Prism 6. Comparisons of significance between groups were performed using unpaired Student *t*-test by Quickcalc.

## 3. Results

### 3.1. Enrichment of lipid droplet small molecular weight proteins and identification of the proteins encoded by noncoding RNAs

Previously we established a method for isolation and proteomic study of LDs and collectively identified many proteins of LDs from bacteria to humans[17; 24]. But the proteins coded from nominally noncoding transcripts have not been identified in LDs. In this study, we used a proteomic-based strategy to discover such proteins in isolated LDs of myoblasts (Fig. 1A). Previously, we constructed a database of proteins derived from noncoding transcripts with sORF, with which we were able to identify 54 novel proteins encoded by noncoding RNAs in mouse serum including 2 of which were further verified to express *in vivo*[25]. In brief, mouse GENCODE (vM4) and Ensembl transcripts (release 73) were used to annotate noncoding transcripts. Then the resulting noncoding transcripts were translated to small ORF-encoded proteins, ranging from 8-to 100-amino acids, by the ORF Finder program and an in-house program. This database, containing a total of 525,792 putative ORFs, was then merged with mouse Uniprot database and Contamination database. The combined database was then trained and verified by current known sORF-encoded proteins. Mass spectrometric data generated in this study were searched against this database.

**Figure 1.**
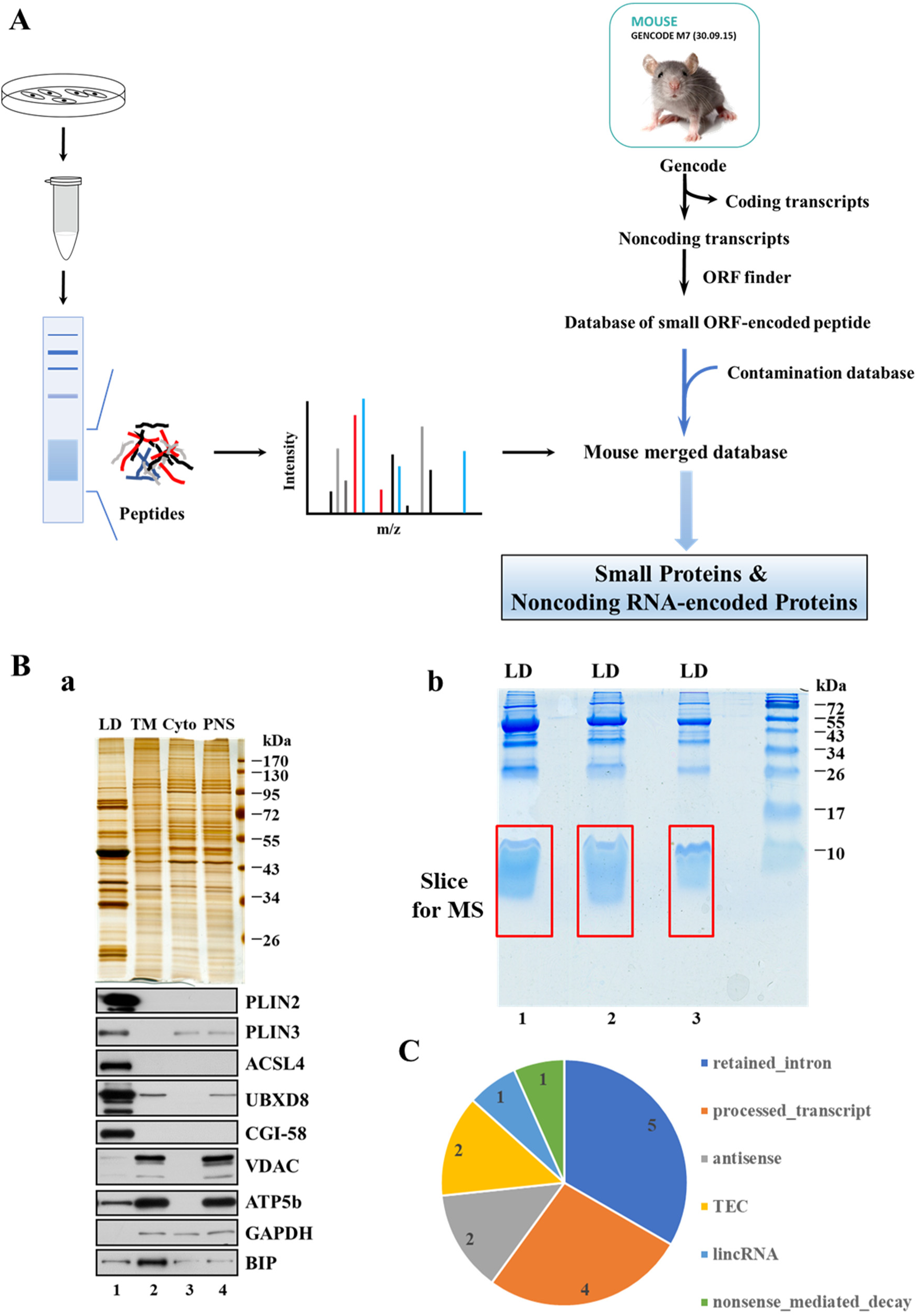
Identification of noncoding RNA-encoded proteins from C2C12 lipid droplets. **A** Workflow for the study. After isolation of LDs, low molecular weight proteins were enriched, digested, and the peptides were subjected to mass spectrometry analysis. The mass spectra obtained were searched against our database, constructed as previously reported. The flowchart for construction of database is adopted and modified from Figure 1 in our previous work[25]. **B a** Quality verification of the isolated LDs. The quality of the isolated LDs from C2C12 was verified by silver staining and Western blot analysis. After isolation, the LD proteins were extracted and subjected to silver staining. Meanwhile, the marker proteins of LD (PLIN2, PLIN3, ACSL4, UBXD8 and CGI-58), endoplasmic reticulum (BIP), mitochondrion (VDAC, ATP5b), and cytosol (GAPDH) were analyzed by Western blot. TM, total membrane; Cyto, cytosol; PNS, post-nuclear supernatant. **b** Enrichment of LD proteins with a molecular mass less than 17 kDa. LD proteins were separated using a 12% Bis-Tris gel and the proteins below 17 kDa were excised for analysis by mass spectrometry. **C** Classification of LD-associated proteins derived from noncoding RNAs. The 15 putative LD-associated proteins encoded by noncoding RNAs were grouped according to transcript of origin. TEC, to be experimentally confirmed.

LDs were isolated from C2C12 myoblasts using our well-established method[23; 24]. The quality of the isolated LD preparation was assessed using silver staining and Western blot methods. In the silver stained gels the protein composition of the isolated LDs are clearly distinct from other cellular fractions including total membrane (TM), cytosol (Cyto), and post-nuclear supernatant (PNS) (Fig. 1Ba, upper panel). The Western blots demonstrate the enrichment of LD marker proteins including PLIN2, PLIN3, ACSL4, UBXD8, and CGI-58 on the isolated LDs (Fig. 1Ba, lower panel). The cytosolic protein GAPDH was not detectable in the LD fraction and the endoplasmic reticulum protein, BIP, was mainly enriched in the membrane fraction. Some mitochondrial proteins were found on LDs, which is consistent with our previous result that there exist LD-anchored mitochondria (LDAM) in skeletal muscle cells[23; 27]. The myoblast LD preparation is of high quality and suitable for further analysis.

Samples from the isolated LDs were applied to 12% Bis-Tris gel to separate and enrich proteins with a molecular weight less than 17 kDa and the gel was stained with Colloidal Blue (Fig. 1Bb). We carried out the isolation of LDs 4 times and those LDs were used for the enrichment and the subsequent proteomic study using our database respectively.

The similarity between three independent LD fractions showed the reproducibility of our system to isolate and enrich small proteins (Fig. 1Bb). The large stained region below 17 kDa contained small proteins and degraded proteins. This was the first study to specifically analyze this population in detail. Among the proteins identified were 136 coded for by classical genes. Within this group, 44 were identified with two or more unique peptides and these are reported in Supplemental Table S1 by their sub-cellular location or function. Within this group 52% have been reported in previously published LD proteomes or have been localized to LDs by other techniques, demonstrating the reliability of our system to purify LD and enrich small proteins as well as the proteomic study. Among the identified proteins, the most abundant were ribosomal (around 23%, Supplemental Fig. S1). Mitochondrial proteins constituted about 20% of the proteins, which is consistent with our previous work. This suggests a tight anchoring between LDs and mitochondria in the oxidative tissues[27]. Another major group consisted of proteins involved in signaling pathways (up to 14%), which is consistent with the hypothesis that LDs function in signal transduction[17]. 7% were associated with protein degradation in the ubiquitin-proteasome system, suggesting a role for LDs in protein degradation, as has been suggested in previous reports[17].

More importantly, the analysis identified 15 proteins encoded by noncoding RNAs and all of them are noncoding RNA encoded-proteins that have not been identified previously. They arose from six transcript classes: retained intron, processed transcript, antisense, TEC (to be experimentally confirmed), lincRNA, and nonsense mediated decay (Fig. 1C). Among those peptides, those arose from retained intron and processed transcript occupied 60%, which indicated that LD-associated micropeptides derived from those two classes may be more than from other transcript classes annotated to be noncoding. Interestingly, a transcript category, TEC (to be experimentally confirmed), which has been specifically created to highlight the coding potential of region that require experimentally validation, was found to code two of the identified peptides.

The putative proteins were named LD-associated noncoding RNA-encoded proteins (LDANPs) and are listed in Table 1 in descending order of their XCorr scores. The XCorr score is an important search algorithm used to assess the quality of a peptide-spectrum match, which indicates the peptide-identification reliability in mass spectrometry.

**Table 1.**
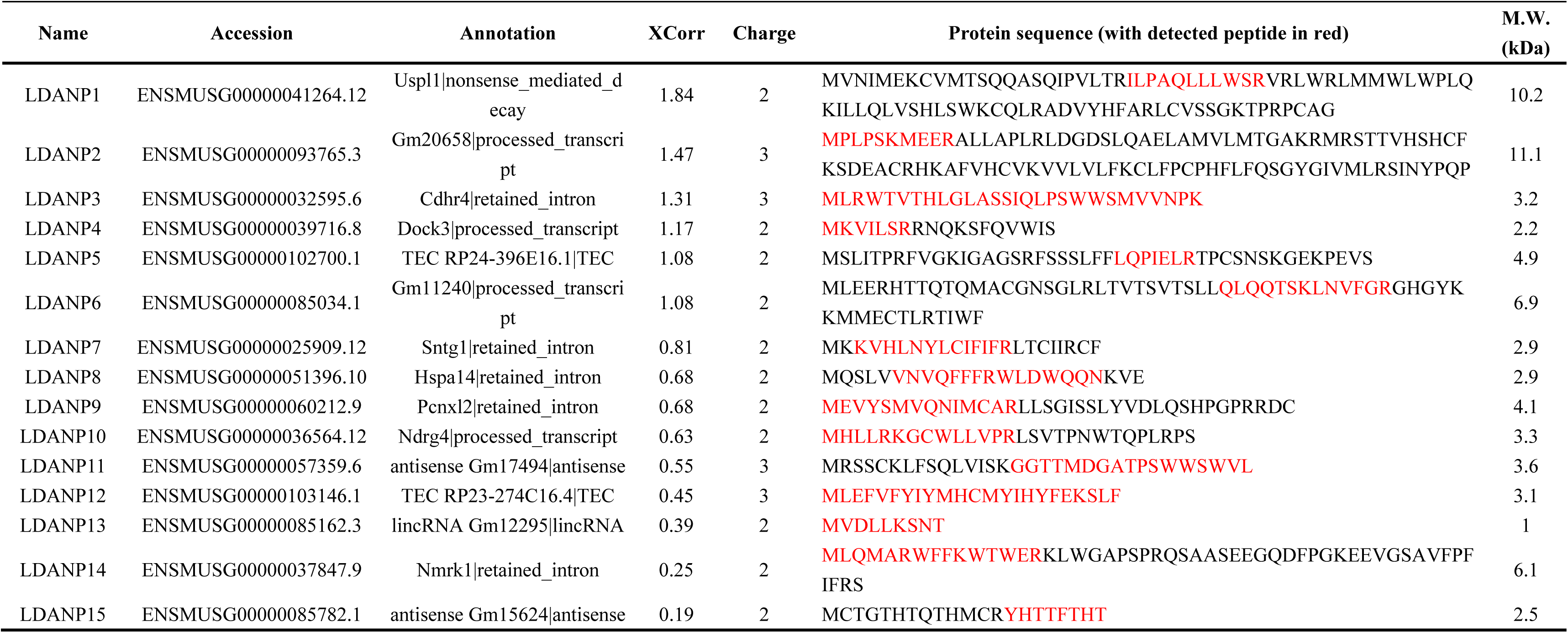
Proteins encoded by nominally noncoding transcripts on C2C12 lipid droplets.

### 3.2. Degradation of LDANP1 is regulated by the proteasome

The protein with the highest XCorr score, LDANP1, was chosen for further study (Table 1). The murine *LDANP1* is located to the exon 4 of ubiquitin specific peptidase-like 1 (*USPL1*) on chromosome 5, containing 261 nucleotides (starting from 149,194,024 to 149,914,284) (Fig. 2A). However, it is translated in the +1 reading frame, resulting in the 87-amino acid protein, LDANP1. To gain insights into the regulation of LDANP1, this protein was overexpressed as a GFP-FLAG fusion protein. It is known that proteins encoded by noncoding RNAs are always expressed in low abundance and are unstable[7]. Besides, LD resident proteins PLIN1 and PLIN2 are degraded through proteasome-dependent pathway[28; 29]. As described above, LDANP1 was identified in isolated LD fraction derived from noncoding transcripts. We wanted to determine if LDANP1 was also degraded by the proteasome.

**Figure 2.**
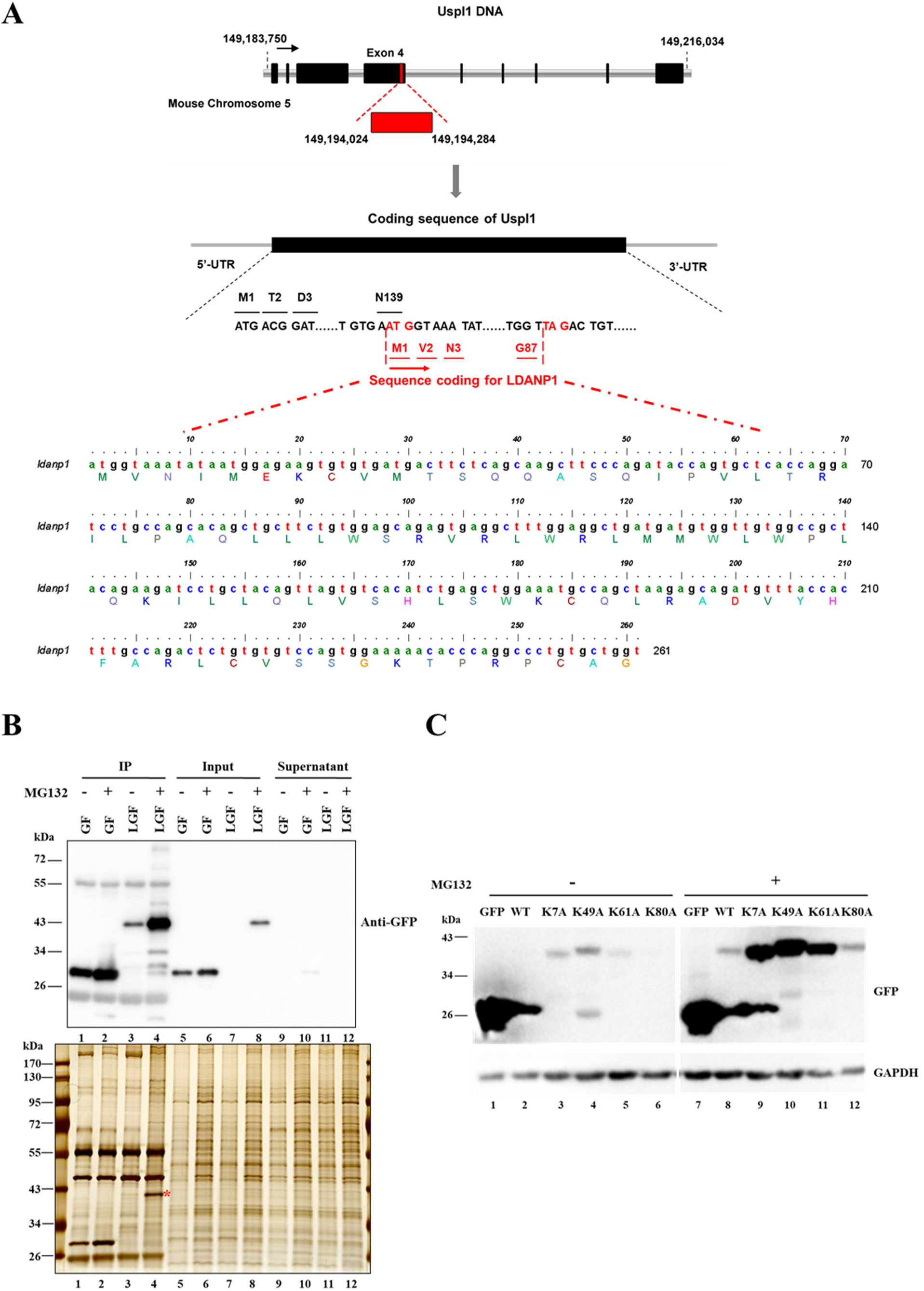
LDANP1 is degraded by the proteasome. **A** Schematic representation of the locus and translation of LDANP1. In mice, *LDANP1* (red) is located on exon 4 of the *USPL1* gene (black) on chromosome 5. LDANP1 is translated in the +1 frame. The nucleotide and amino acid sequences are shown (using BioEdit). **B** Effect of MG132 on the expression of LDANP1. Cells stably expressing LDANP1-GFP-FLAG (LGF) and GFP-FLAG control cells (GF) were treated with or without 1 µM MG132 for 12 h. Cells were washed with PBS, lysed with 1 mL TETN solution and probed with FLAG beads to precipitate FLAG tagged protein. Precipitate was eluted in sample buffer and separated on 10% SDS-PAGE. The silver stained gel (lower panel) displayed the protein profile in each sample and served as protein loading control. Separated proteins were blotted and probed with anti-GFP antibody (upper panel). GF, GFP-FLAG cells; LGF, LDANP1-GFP-FLAG cells; IP, immunoprecipitation. **C** Analysis of potential ubiquitin acceptor site(s) on LDANP1. Lysine residues in LDANP1 were mutated to alanine using LDANP1-GFP as template. Cells stably expressing mutated LDANP1 and control cells were treated with or without 1 µM MG132 for 12 h. Whole cell lysate were probed with GFP and GAPDH, the loading control. WT, cells expressing LDANP1-GFP.

Control cells expressing GFP-FLAG (GF) and cells stably expressing LDANP1-GFP-FLAG (LGF) were cultured in media with and without MG132, a specific proteasome inhibitor. FLAG immunoprecipitation was performed followed by GFP Western blot. After addition of MG132 a band of approximately 40 kDa, consistent with the molecular weight of LDANP1 fused with GFP and FLAG, was detected in whole cell lysate of the LGF cells (Fig. 2B, upper panel, lane 8). MG132 treatment also resulted in a dramatic increase in the intensity of a 40 kDa band in the immunoprecipitate from the LGF cells by GFP Western blot (Fig. 2B, upper panel, lane 4) and silver staining (Fig. 2B, lower panel, lane 4, red star). These results demonstrate that LDANP1 was degraded through the proteasome.

To identify the potential ubiquitin acceptor site(s) on LDANP1, all four lysine residues were mutated to alanine independently. Then, the respective contribution of each residue to peptide stability was evaluated (Fig. 2C). LDANP1 and LDANP1 mutants, all fused with GFP, were stably expressed in C2C12 cells and were evaluated for protein level by Western blot. Lysine residues 7, 49, and 61 influenced LDANP1 stability, as shown by the increase of GFP signal in cell lysates, compared to cells expressing WT LDANP1-GFP (Fig. 2C, lanes 3-5 vs. lane 2). Meanwhile, the lysine 80 mutant appeared to be unstable and the GFP signal was not detectable in cells not treated with MG132, similar to WT (Fig. 2C, lane 6 vs. lane 2). Therefore, lysine residue 80 is most likely not ubiquitinated. However, when treated with MG132, all GFP signals were detected (Fig. 2C, lanes 9-12). Taken together, these results demonstrate that LDANP1 is regulated by proteasome protein degradation and suggest that multiple lysine residues on LDANP1 are potential sites of regulation by the ubiquitin system.

### 3.3. LDANP1 is localized on lipid droplets in C2C12 cells

LDANP1 was identified in the LD proteome and was found to be degraded in a proteasome-dependent manner like LD resident proteins. We next set out to confirm its localization on LDs using biochemical and morphological methods. First, we transiently expressed the protein with GFP tag on the C terminus. Figure 3A shows clear GFP fluorescence in ring structures around LDs stained by LipidTOX Red, indicating that it was co-localized with LDs.

**Figure 3.**
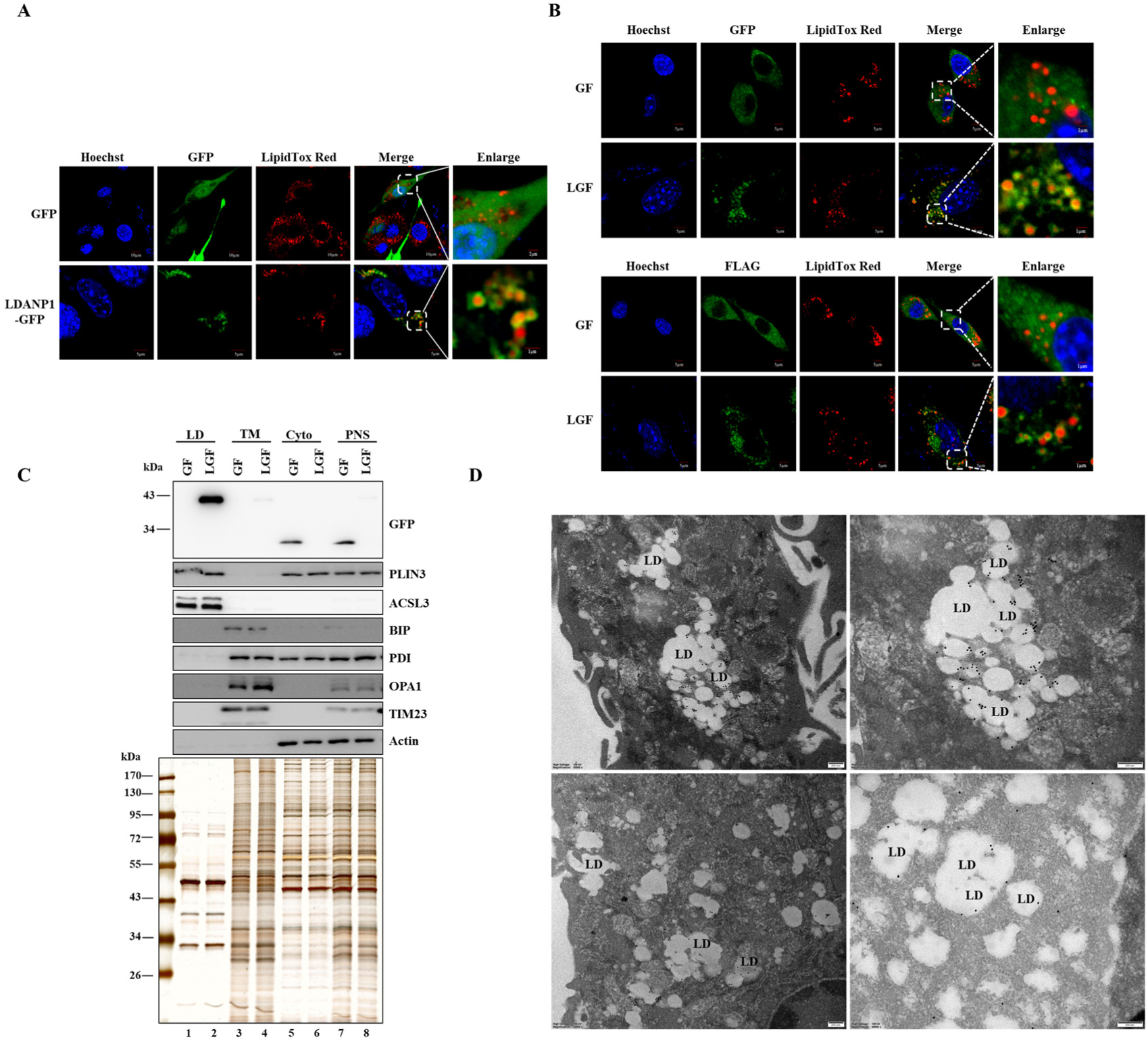
LDANP1 localizes to lipid droplets in C2C12 cells. The localization of LDANP1 in C2C12 cells was studied using several methods. **A** The localization of transiently expressed-LDANP1. LDANP1-GFP was transiently expressed in C2C12 cells and treated with 100 µM OA for 12 h. Twenty-four hours post-transfection, cells were stained with LipidTOX Red and Hoechst (blue) to stain LDs and the nucleus, respectively. The cells were viewed using a confocal microscope. In the upper panel, Bar = 10 μm, except for Bar = 2 μm in the enlarged image. In lower panel, Bar = 5 μm, except for Bar = 1 μm in the enlarged image. **B** The localization of stably expressed-LDANP1. C2C12 cells stably expressing GFP-FLAG and LDANP1-GFP-FLAG were treated with 100 µM OA for 12 h. Then the cells were analyzed by immunofluorescence staining for GFP or FLAG with green-fluorescent signal to show the localization of stably expressed-LDANP1. Subsequently, the cells were stained with LipidTOX Red for LDs and Hoechst for nuclei. The cells were viewed using a confocal microscope. Bar = 5 μm, except for Bar = 1 μm in the enlarged image. **C** Localization of LDANP1 demonstrated by cell fractionation. C2C12 cells stably expressing LDANP1-GFP-FLAG and GFP-FLAG control cells were cultured in 100 µM OA supplemented medium for 12 h. Cells were harvested for cell fractionation. The proteins in different cellular fractions were separated by SDS-PAGE and analyzed by Western blot with the indicated antibodies. GF, GFP-FLAG cells; LGF, LDANP1-GFP-FLAG cells. TM, total membrane; Cyto, cytosol; PNS, post-nuclear supernatant. **D** Immuno-gold labeling for LDANP1. C2C12 cells stably expressing LDANP1-GFP-FLAG were treated with 100 µM OA plus 1 µM MG132 for 12 h. Then the cells were prepared for immuno-gold labeling of GFP to display the localization of LDANP1. Briefly, cells were fixed with 4% (v/v) paraformaldehyde, dehydrated in an ascending concentration series of ethanol and embedded in LR White resin. After ultra-thin sectioning, sections were stained for GFP with secondary antibody conjugated to 18 nm colloidal gold. The sections were observed by transmission electron microscope. Bar = 200 nm.

The localization of LDANP1 in the LDANP1-GFP-FLAG stable cells was confirmed by immunofluorescence staining of GFP or FLAG (green signal, Fig. 3B). Consistent with the transiently transfected cells, the LDANP1 was also localized to LDs in stable cells.

The localization of LDANP1 was accessed through cell fractionation. C2C12 cells stably expressing LDANP1-GFP-FLAG was fractionated by differential centrifugation and the fractions were analyzed by Western blotting with anti-GFP. The LDANP1-GFP-FLAG co-fractionated with known LD-associated proteins, PLIN3 and ACSL3 (Fig. 3C, upper panel, lane 2). In control cells, GFP-FLAG co-fractioned with cytosolic protein Actin (Fig. 3C, upper panel, lane 5). Minimal GFP signal could be detected in the membrane fraction, which contained endoplasmic reticulum, as marked by BIP and PDI, and mitochondrion, indicated by OPA1 and TIM23 (Fig. 3C, upper panel, lane 4). This biochemical result also revealed that LDANP1 was heavily enriched in LD fraction.

Finally, LDANP1 localization was also determined using immuno-gold labeling. Figure 3D shows that the 18-nm colloidal gold particles labeling GFP in LDANP1-GFP-FLAG stable cells mainly surrounded LDs (black dots).

All lines of evidence clearly demonstrate that LDANP1 localizes to LDs in C2C12 cells.

### 3.4. LDANP1 is endogenously expressed in C2C12 cells

The next essential question to be addressed was whether LDANP1 is expressed endogenously. First, LDANP1 was analyzed computationally for conservation. Alignment of the LDANP1 amino acid sequence against in silico-translated short ORFs in USPL1 sequences from 11 different mammalian species (Fig. 4A) revealed that this protein is conserved and 48% of amino acid residues appeared to be invariant between human and mouse. Therefore, the protein is broadly conserved across mammals. Second, mass spectrometry detected peptide derived from LDANP1 (listed in red in Table 1). Third, a database for analysis of alternatively translated proteins in human RefSeq transcripts also identified a LDANP1 homolog in human within the USPL1 coding sequence and later also in several other mammals[30]. These data suggested the possibility of endogenous translation of LDANP1. Therefore, we used CRISPR (Clustered Regularly Interspaced Short Palindromic Repeats)-Cas9 (CRISPR-associated protein 9)-mediated gene editing strategy to validate its endogenous expression. A 3×FLAG-HA sequence was integrated at the C-terminus of and in-frame with LDANP1 ORF located within USPL1 exon 4 in C2C12 cells (Fig. 4B).

**Figure 4.**
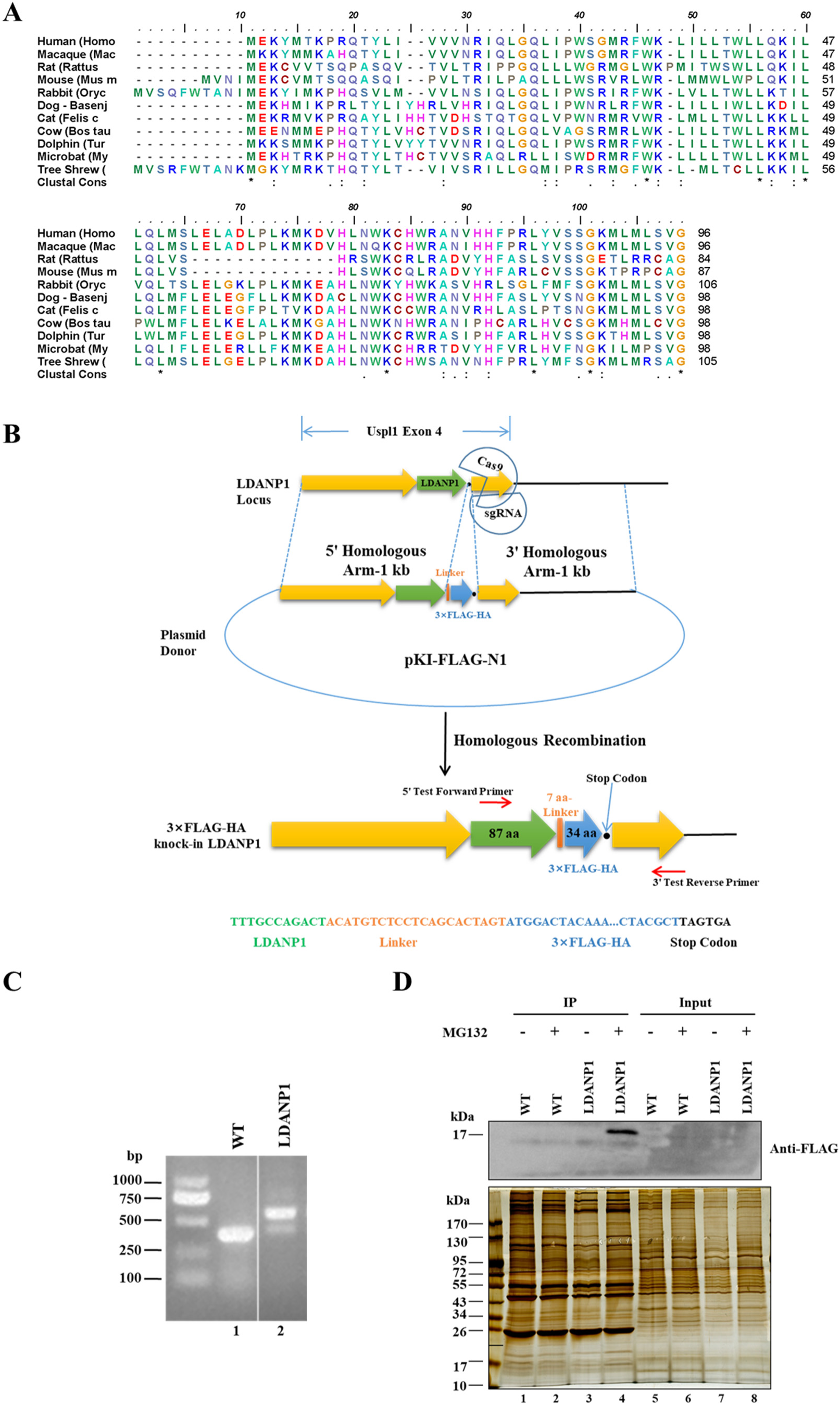
Verification of endogenous LDANP1 expression. **A** Alignment of putative LDANP1 peptide sequences from a variety of mammals. LDANP1 amino acid sequences were translated from short ORF in USPL1 sequence in 11 different mammalian species. Alignment of those sequences were analyzed using BioEdit. Amino acid identity is indicated. **B** Schematic depicting CRISPR/Cas9-mediated LDANP1 knock-in strategy. Cells carrying a 3×FLAG-HA tag in the C-terminus of LDANP1 allele were generated by CRISPR/Cas9-mediated knock-in technique. **C** Verification of the insertion of the 3×FLAG-HA tag by PCR. Genomic DNA from selected clones was probed with primer pairs designed to amplify 200 bp upstream and 200 bp downstream of the LDANP1 stop codon. The PCR amplicons were subjected to 2% (w/v) agarose gel electrophoresis. **D** Verification of endogenous translation of LDANP1. LDANP1 knock-in cell line and WT C2C12 cells were treated with or without 1 µM MG132 for 12 h. Cells were washed with PBS, lysed with 1 mL TETN solution and probed with anti-FLAG beads to precipitate FLAG-tagged protein. The precipitates were eluted in sample buffer, separated on 5-15% gradient SDS-PAGE, and analyzed by FLAG Western blot (upper panel). A silver-stained gel (lower panel) indicated the protein profile in each sample and served as protein loading control. IP, immunoprecipitation.

Correct insertion of the 3×FLAG-HA (123 bp) was verified by polymerase chain reaction (PCR) using a pair of primers targeting nucleotide sequence upstream and downstream of LDANP1 stop codon. Agarose gel electrophoresis of the amplicon from the target genomic region indicated the presence of two bands in LDANP1 knock-in cells (Fig. 4C, lane 2). The lower band, adjacent to the WT, corresponded to the endogenous non-edited allele and the upper band corresponded precisely to edited allele containing 3×FLAG-HA, indicating endogenous expression at the mRNA level. Endogenous expression at the protein level was verified subsequently by Western blot. Since the stability of overexpressed LDANP1 is regulated by the proteasome as previously shown (Fig. 2), the 3×FLAG-HA tagged LDANP1 knock-in C2C12 cells were cultured with or without 1 µM MG132 for 12 h. Whole cell lysate from WT and LDANP1 knock-in cell lines were subjected to FLAG immunoprecipitation to concentrate the protein. The FLAG immunoprecipitation sample was further analyzed with anti-FLAG and the result showed a distinct band of approximately 17 kDa in the MG132-treated LDANP1 knock-in cells (Fig. 4D, upper panel, lane 4). This unique band lies in the predicted molecular weight region for the 3×FLAG-HA knock-in LDANP1 (10.2 kDa of LDANP1 and 4.8 kDa of 3×FLAG and HA tag, encoded in 34 amino acids, plus a 7-amino acid linker). Therefore, LDANP1 is translated endogenously.

### 3.5. LDANP1 reduces triacylglycerol storage and mediates insulin sensitivity in C2C12 cells

LDANP1 is the first protein derived from a nominally noncoding transcript found on LDs. To examine its biological function on LD dynamics, C2C12 cells stably expressing GFP-FLAG and LDANP1-GFP-FLAG were incubated in the presence or absence of 100 µM oleate (OA) for 12 h. Total triacylglycerol (TAG) levels in GFP-FLAG control and LDANP1-GFP-FLAG cell lines treated with vehicle (Fig. 5A) showed no significant differences. However, the TAG level was significantly lower in the LDANP1 stably expressed cells treated with OA compared to the GFP-FLAG control. This suggests that LDANP1 negatively regulates OA-mediated TAG storage. OA is mainly incorporated into TAG in C2C12 cells and has been shown by our group and others to reverse the inhibitory effect of saturated fatty acids on insulin signaling by promoting their incorporation into TAG[22; 31; 32]. Therefore, we wondered whether LDANP1 affects insulin signaling.

**Figure 5.**
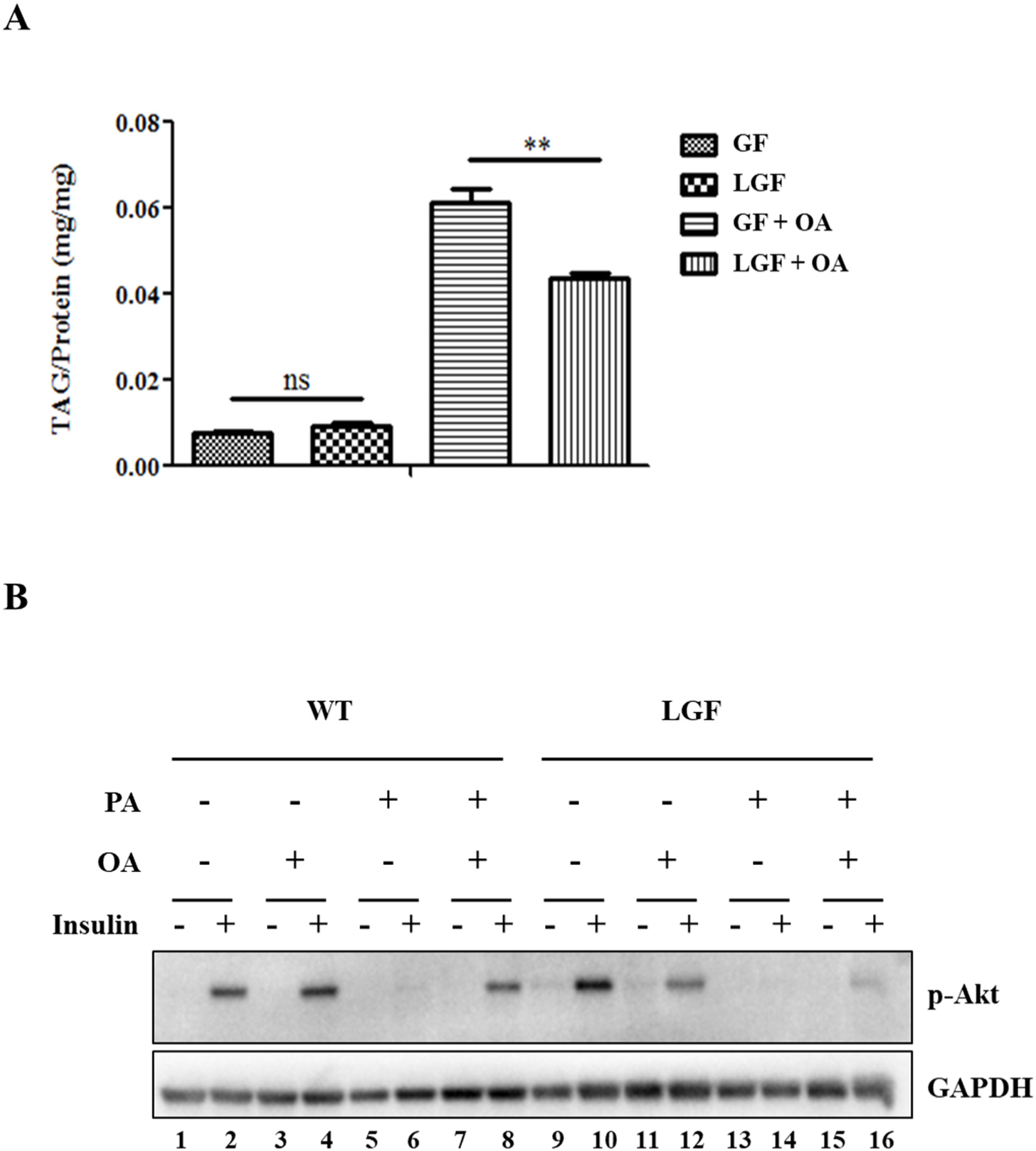
LDANP1 reduces the storage of triacylglycerol and mediates insulin sensitivity in C2C12 cells. **A** Reduction of triacylglycerol level by LDANP1. Stable cells expressing LDANP1-GFP-FLAG and GFP-FLAG control cells were cultured to confluence in 6-well dishes and treated with or without 100 µM oleate (OA) for 12 h. Cells were collected in PBS (pH 7.4) containing 1% (v/v) Triton X-100 and sonicated. An aliquot of cell lysate was analyzed for triacylglycerol (TAG) using total protein as internal control. Data represent mean ± SEM (n = 4), and were analyzed by unpaired Student *t*-test. **, P<0.001. GF, GFP-FLAG cells; LGF, LDANP1-GFP-FLAG cells. **B** Inhibitory effect of LDANP1 on insulin sensitivity in C2C12 cells. Cells were incubated in the presence or absence of 100 μM OA with or without 500 μM palmitate (PA) for 12 h before insulin stimulation (200 nM insulin for 10 min at 37°C). Cells were then lysed with sample buffer and the proteins were subjected to Western blot analysis of p-Akt and GAPDH. LGF, LDANP1-GFP-FLAG cells.

In agreement with our previous report, PA treatment reduced insulin signal represented by Akt phosphorylation (p-Akt) in WT C2C12 cells (Fig. 5B, lane 6 vs lane 2). Treatment with OA alone had no significant effect (Fig. 5B, lane 4 vs lane 2). However, when the WT cells were incubated with both PA and OA, the PA-suppressed p-Akt signal was recovered (Fig. 5B, lane 8 vs lanes 6 and 2). In LDANP1 stably expressed cells, PA still reduced the p-Akt signal similarly to WT (Fig. 5B, lane 14 vs lane 10), but OA could not rescue PA-suppressed insulin signaling (Fig. 5B, lane 16 vs lane 10). Therefore, LDANP1 functions to reduce the ability of TAG storage and, in turn, influences insulin sensitivity in C2C12 cells.

## 4. Discussion

Much progress has been made in recent years in identifying the full set of sequences that can be translated in cells, a complement which extends beyond the classically recognized long proteins. Many noncoding sequences have been identified to habor small ORFs that potentially encode functional proteins, micropeptides, less than 100 amino acids in length. Some of those proteins have been found to play significant biological roles in various cellular processes. The LD, the prominent reservoir for lipid energy and membrane building blocks in cells, plays a crucial role in lipid metabolism and storage, and is actively involved in metabolic diseases. No proteins encoded by noncoding sORFs have been reported on LDs.

We aimed to determine if there are proteins encoded by nominally noncoding RNAs on LDs. A bottom-up strategy is adopted here, that is, the study starts with the isolated organelle directly other than from the whole cell. Using mass spectrometry we identified 15 novel proteins, derived from nominally noncoding transcripts, on LDs isolated from C2C12 cells and validated the endogenous expression of one of them. This is the first report to study the proteins from noncoding sORFs on LDs directly and this strategy may be applicable in searching micropeptides from other cellular organelles.

This is the first systematic study of LD-associated proteins encoded by noncoding sORFs. Several traits are apparent from these micropeptides identified in isolated LDs. First, the distribution pattern of those peptides (Fig. 1C) across different transcript classes indicates that the micropeptides associated with LDs may be more common in retained intron and processed transcript. Second, those micropeptides are present in very low abundance. Unlike the classically translated proteins, such as ribosome proteins, half of which were detected more than 3 fragments in mass spectrometry, all micropeptides were matched with only one fragment (Table 1). Even a stably expressed, recombinant LDANP1 could not be detected in the whole cell lysate (Fig. 2B, upper panel, lane 7). The protein is only detectable after enrichment by immunoprecipitation (Fig. 2B, upper panel, lane 3).

In this study, LDANP1 was chosen for further study due to its high XCorr score. This protein was verified to localize on LDs in C2C12 cells through several experimental techniques. The endogenous expression of the protein was verified using a CRISPR/Cas9-mediated gene editing approach. The results demonstrate that this is an authentic and novel small protein found on LDs. More than 100 proteins have been reported to be associated with LDs. LDs have been recognized as an important organelle for only the last two decades and many fundamental questions in LD biology remain unanswered[33]. Therefore, the presence of proteins derived from nominally noncoding transcripts represents a new frontier in study of LDs.

Interestingly, the degradation of LDANP1 was found to be proteasome-dependent, similar to the degradation of LD resident proteins, PLIN1 and PLIN2[28; 29]. This supports the proposal that proteasomal regulation is a common mechanism for the modulation of LD-associated proteins[29]. We identified multiple lysine residues on LDANP1 important for ubiquitination in the proteasome system. However, ubiquitination plays other important roles besides targeting proteins for degradation, including DNA repair, signal transduction, and endocytosis[34]. Therefore, it remains unclear whether the multiple ubiquitination sites of LDANP1 serve other purposes besides driving degradation or if such a signature is common to proteins derived from small ORFs to facilitate their rapid turnover.

No specific LD targeting signal has been found but LD-associated proteins that directly target to LDs usually contain a hydrophobic domain or amphipathic α-helix[17; 18]. For instance, LD resident protein in plants, oleosin, is comprised of an N-terminal amphipathic domain, a central hydrophobic domain, and a C-terminal amphipathic α-helical domain. The central domain anchors oleosin into the hydrophobic core of LD and the N-and C-terminal domains are proposed to reside on the monolayer phospholipid membrane of LDs[35; 36]. Using bioinformatic tools, we analyzed the possible secondary structure of LDANP1 (Supplemental Information). LDANP1 was predicted to have one amphipathic α-helix on each terminus, spanned by a hydrophobic α-helix. We propose that LDANP1 targets LDs in a manner similar to oleosin, with the hydrophobic helix anchoring LDANP1 in LD hydrophobic core and two amphipathic α-helices binding the monolayer phospholipid membrane of LDs (Supplemental Fig. S2). We are pursuing further investigation into its exact targeting mechanism.

LDANP1 is identified as an alternative protein derived as an out-of-frame ORF in the USPL1 coding sequence. Previous work suggests that dual coding is more prevalent in genes with low expression levels because genes which are highly expressed are optimized for efficient translation and their coding sequences are too restrained to accommodate additional coding information[37]. Previous work on USPL1 indicated that the protein is translated in low abundance in all cell lines examined[38], which further supports the fact that genes encoding low abundant proteins might be a reservoir of alternative proteins. Interestingly, while USPL1 is located to the nucleus and affects coilin localization[38], LDANP1 is shown to localize on LDs and influence lipid metabolism, suggesting a variety of functions for alternatively translated proteins.

Skeletal muscle consumes the majority of glucose in humans and plays a crucial role in maintaining glucose homeostasis[39]. Excessive lipid storage in myocytes is closely related to insulin resistance[19]. Determining the mechanism linking TAG storage and insulin sensitivity in skeletal muscle is an essential goal in the study of insulin resistance. In our and others previous studies, PA was found to inhibit insulin sensitivity through a conversion into active lipid molecules. OA can protect insulin signaling from the inhibitory effects of PA in myoblasts by driving the fatty acids into TAG[22; 31; 32]. As a LD-associated protein, LDANP1 was found to reduce the storage of TAG when myocytes were loaded with OA. In turn, the rescue of PA-reduced insulin sensitivity by OA was suppressed in the presence of an excess of LDANP1. Therefore, we propose that this protein could block the incorporation of OA into TAG thus affecting insulin sensitivity in myoblast. However, the mechanism is unknown. One should work on identifying its potential interaction partner(s), since several known micropeptides are found to exert functions by engaging with and modulating larger regulatory proteins[7]. Meanwhile, its effect on insulin sensitivity in the skeletal muscle of mice will also be investigated. Taken together, this study sheds light on novel factors, LD-associated micropeptides and/or their functional machineries, which can affect LD dynamics and insulin signaling regulation in skeletal muscle.

## 5. Conclusions

In this study, we have shown for the first time proteins derived from nominally noncoding transcripts on LDs. The localization of one of these proteins, LDANP1, was verified to localize on LDs by multiple experimental techniques. This protein reduces the TAG storage and the storage-related insulin sensitivity in myoblasts. Those findings suggest the vast pool of putative proteins derived from nominally noncoding transcripts serves as candidates for screening new LD-associated proteins as well as factors modulating LD dynamics and lipid metabolism.

## Abbreviations

LD: lipid droplet
LDANP: LD-associated noncoding RNA-encoded proteins
sORFs: small open reading frames
OA: oleate
TAG: triacylglycerol
PA: palmitate
TM: total membrane
Cyto: cytosol
PNS: post-nuclear supernatant
LDAM: LD-anchored mitochondria
GF: cells expressing GFP-FLAG
LGF: cells stably expressing LDANP1-GFP-FLAG
CRISPR/Cas9: clustered regularly interspaced short palindromic repeats/CRISPR-associated protein 9

## Author contributions

S.Z. and P.L. designed the project. T.H., A.B., and S.X. performed the experiments. Y.D. helped with the immunoprecipitation analysis. K.X. and O.O. helped with triacylglycerol assay. A.H.M. helped with bioinformatic analysis. J.W. helped with MS study. S.Z., P.L., and A.B. wrote the manuscript. All authors have read and approved the manuscript.

## Acknowledgments

The authors thank Dr. John Zehmer for his critical reading and useful suggestions. The authors also thank Mr. Lianwan Chen for his help in immunogold labeling experiment and Ms. Chen Ye for her critical reading. The authors also thank Ms. Yan Teng for her help in confocal imaging. This work was supported by the National Key R&D Program of China (Grant No. 2018YFA0800900, 2016YFA0500100 and 2018YFA0800700), National Natural Science Foundation of China (Grant No. 31671402, 91954108, 91857201, 31671233, 31701018 and U1702288). This work was also supported by the “Personalized Medicines——Molecular Signature-based Drug Discovery and Development”, Strategic Priority Research Program of the Chinese Academy of Sciences, Grant No. XDA12040218.

## Competing financial interests

The authors declare no competing financial interests.

## Notes

### Competing Interest Statement

The authors have declared no competing interest.

## References

[1] Lander, E.S., 2011. Initial impact of the sequencing of the human genome. Nature 470(7333):187–197.

[2] Carninci, P., Kasukawa, T., Katayama, S., Gough, J., Frith, M.C., Maeda, N., et al., 2005. The transcriptional landscape of the mammalian genome. Science 309(5740):1559–1563.

[3] Kapranov, P., Cheng, J., Dike, S., Nix, D.A., Duttagupta, R., Willingham, A.T., et al., 2007. RNA maps reveal new RNA classes and a possible function for pervasive transcription. Science 316(5830):1484–1488.

[4] Ruiz-Orera, J., Alba, M.M., 2019. Translation of Small Open Reading Frames: Roles in Regulation and Evolutionary Innovation. Trends in Genetics 35(3):186–198.

[5] Consortium, E.P., 2012. An integrated encyclopedia of DNA elements in the human genome. Nature 489(7414):57–74.

[6] Hon, C.C., Ramilowski, J.A., Harshbarger, J., Bertin, N., Rackham, O.J., Gough, J., et al., 2017. An atlas of human long non-coding RNAs with accurate 5’ ends. Nature 543(7644):199–204.

[7] Makarewich, C.A., Olson, E.N., 2017. Mining for Micropeptides. Trends in Cell Biology 27(9):685–696.

[8] Sousa, M.E., Farkas, M.H., 2018. Micropeptide. PLoS Genetics 14(12):e1007764.

[9] Pauli, A., Norris, M.L., Valen, E., Chew, G.L., Gagnon, J.A., Zimmerman, S., et al., 2014. Toddler: an embryonic signal that promotes cell movement via Apelin receptors. Science 343(6172):1248636.

[10] Anderson, D.M., Anderson, K.M., Chang, C.L., Makarewich, C.A., Nelson, B.R., McAnally, J.R., et al., 2015. A micropeptide encoded by a putative long noncoding RNA regulates muscle performance. Cell 160(4):595–606.

[11] Lee, C., Zeng, J., Drew, B.G., Sallam, T., Martin-Montalvo, A., Wan, J., et al., 2015. The mitochondrial-derived peptide MOTS-c promotes metabolic homeostasis and reduces obesity and insulin resistance. Cell Metabolism 21(3):443–454.

[12] Matsumoto, A., Pasut, A., Matsumoto, M., Yamashita, R., Fung, J., Monteleone, E., et al., 2017. mTORC1 and muscle regeneration are regulated by the LINC00961-encoded SPAR polypeptide. Nature 541(7636):228–232.

[13] Martin, S., Parton, R.G., 2006. Lipid droplets: a unified view of a dynamic organelle. Nature Reviews. Molecular Cell Biology 7(5):373–378.

[14] Zhang, C., Liu, P., 2017. The lipid droplet: A conserved cellular organelle. Protein and Cell 8(11):796–800.

[15] Zehmer, J.K., Huang, Y., Peng, G., Pu, J., Anderson, R.G., Liu, P., 2009. A role for lipid droplets in inter-membrane lipid traffic. Proteomics 9(4):914–921.

[16] Farese, R.V., Jr., Walther, T.C., 2009. Lipid droplets finally get a little R-E-S-P-E-C-T. Cell 139(5):855–860.

[17] Zhang, C., Liu, P., 2019. The New Face of the Lipid Droplet: Lipid Droplet Proteins. Proteomics 19(10):e1700223.

[18] Xu, S., Zhang, X., Liu, P., 2018. Lipid droplet proteins and metabolic diseases. Biochimica et Biophysica Acta 1864(5 Pt B):1968–1983.

[19] Samuel, V.T., Shulman, G.I., 2012. Mechanisms for insulin resistance: common threads and missing links. Cell 148(5):852–871.

[20] Crunk, A.E., Monks, J., Murakami, A., Jackman, M., Maclean, P.S., Ladinsky, M., et al., 2013. Dynamic regulation of hepatic lipid droplet properties by diet. PLoS ONE 8(7):e67631.

[21] Bosma, M., Hesselink, M.K., Sparks, L.M., Timmers, S., Ferraz, M.J., Mattijssen, F., et al., 2012. Perilipin 2 improves insulin sensitivity in skeletal muscle despite elevated intramuscular lipid levels. Diabetes 61(11):2679–2690.

[22] Peng, G., Li, L., Liu, Y., Pu, J., Zhang, S., Yu, J., et al., 2011. Oleate blocks palmitate-induced abnormal lipid distribution, endoplasmic reticulum expansion and stress, and insulin resistance in skeletal muscle. Endocrinology 152(6):2206–2218.

[23] Zhang, H., Wang, Y., Li, J., Yu, J., Pu, J., Li, L., et al., 2011. Proteome of skeletal muscle lipid droplet reveals association with mitochondria and apolipoprotein a-I. Journal of Proteome Research 10(10):4757–4768.

[24] Ding, Y., Zhang, S., Yang, L., Na, H., Zhang, P., Zhang, H., et al., 2013. Isolating lipid droplets from multiple species. Nature Protocols 8(1):43–51.

[25] Deng, Y., Bamigbade, A.T., Hammad, M.A., Xu, S., Liu, P., 2018. Identification of small ORF-encoded peptides in mouse serum. Biophys Rep 4(1):39–49.

[26] Wang, Z., 2009. Epitope tagging of endogenous proteins for genome-wide chromatin immunoprecipitation analysis. Methods in Molecular Biology 567:87–98.

[27] Cui, L., Mirza, A.H., Zhang, S., Liang, B., Liu, P., 2019. Lipid droplets and mitochondria are anchored during brown adipocyte differentiation. Protein and Cell 10(12):921–926.

[28] Masuda, Y., Itabe, H., Odaki, M., Hama, K., Fujimoto, Y., Mori, M., et al., 2006. ADRP/adipophilin is degraded through the proteasome-dependent pathway during regression of lipid-storing cells. Journal of Lipid Research 47(1):87–98.

[29] Xu, G., Sztalryd, C., Londos, C., 2006. Degradation of perilipin is mediated through ubiquitination-proteasome pathway. Biochimica et Biophysica Acta 1761(1):83–90.

[30] Vanderperre, B., Lucier, J.F., Roucou, X., 2012. HAltORF: a database of predicted out-of-frame alternative open reading frames in human. Database (Oxford) 2012:bas025.

[31] Yu, J., Li, Y., Zou, F., Xu, S., Liu, P., 2015. Phosphorylation and function of DGAT1 in skeletal muscle cells. Biophys Rep 1:41–50.

[32] Palomer, X., Pizarro-Delgado, J., Barroso, E., Vazquez-Carrera, M., 2018. Palmitic and Oleic Acid: The Yin and Yang of Fatty Acids in Type 2 Diabetes Mellitus. Trends Endocrinol Metab 29(3):178–190.

[33] Welte, M.A., 2015. Expanding roles for lipid droplets. Current Biology 25(11):R470–481.

[34] Passmore, L.A., Barford, D., 2004. Getting into position: the catalytic mechanisms of protein ubiquitylation. Biochemical Journal 379(Pt 3):513–525.

[35] Huang, A.H., 1996. Oleosins and oil bodies in seeds and other organs. Plant Physiology 110(4):1055–1061.

[36] Shao, Q., Liu, X., Su, T., Ma, C., Wang, P., 2019. New Insights Into the Role of Seed Oil Body Proteins in Metabolism and Plant Development. Front Plant Sci 10:1568.

[37] Michel, A.M., Choudhury, K.R., Firth, A.E., Ingolia, N.T., Atkins, J.F., Baranov, P.V., 2012. Observation of dually decoded regions of the human genome using ribosome profiling data. Genome Research 22(11):2219–2229.

[38] Schulz, S., Chachami, G., Kozaczkiewicz, L., Winter, U., Stankovic-Valentin, N., Haas, P., et al., 2012. Ubiquitin-specific protease-like 1 (USPL1) is a SUMO isopeptidase with essential, non-catalytic functions. EMBO Reports 13(10):930–938.

[39] Shulman, G.I., Rothman, D.L., Jue, T., Stein, P., DeFronzo, R.A., Shulman, R.G., 1990. Quantitation of muscle glycogen synthesis in normal subjects and subjects with non-insulin-dependent diabetes by 13C nuclear magnetic resonance spectroscopy. New England Journal of Medicine 322(4):223–228.

